# Probing effects of the SARS-CoV-2 E protein on membrane curvature and intracellular calcium

**DOI:** 10.1101/2021.05.28.446179

**Authors:** Aujan Mehregan, Sergio Pérez-Conesa, Yuxuan Zhuang, Ahmad Elbahnsi, Diletta Pasini, Erik Lindahl, Rebecca J Howard, Chris Ulens, Lucie Delemotte

**Affiliations:** Laboratory of Structural Neurobiology, Department of Cellular and Molecular Medicine, Faculty of Medicine, KU Leuven, Leuven, Belgium; Department of Applied Physics, Science for Life Laboratory, KTH Royal Institute of Technology, Solna, Sweden; Department of Biochemistry and Biophysics, Science for Life Laboratory, Stockholm University, Solna, Sweden

## Abstract

SARS-CoV-2 contains four structural proteins in its genome. These proteins aid in the assembly and budding of new virions at the ER-Golgi intermediate compartment (ERGIC). Current fundamental research efforts largely focus on one of these proteins – the spike (S) protein. Since successful antiviral therapies are likely to target multiple viral components, there is considerable interest in understanding the biophysical role of its other structural proteins, in particular structural membrane proteins. Here, we have focused our efforts on the characterization of the full-length envelope (E) protein from SARS-CoV-2, combining experimental and computational approaches. Recombinant expression of the full-length E protein from SARS-CoV-2 reveals that this membrane protein is capable of independent multimerization, possibly as a tetrameric or smaller species. Fluorescence microscopy shows that the protein localizes intracellularly, and coarse-grained MD simulations indicate it causes bending of the surrounding lipid bilayer, corroborating a potential role for the E protein in viral budding. Although we did not find robust electrophysiological evidence of ion-channel activity, cells transfected with the E protein exhibited reduced intracellular Ca^2+^, which may further promote viral replication. However, our atomistic MD simulations revealed that previous NMR structures are relatively unstable, and result in models incapable of ion conduction. Our study highlights the importance of using high-resolution structural data obtained from a full-length protein to gain detailed molecular insights, and eventually permitting virtual drug screening.

## Introduction

On March 11, 2020, the World Health Organization (WHO) declared the SARS-CoV-2 outbreak a global pandemic. The scale and the severity of the disease, as well as the speed at which this virus is still spreading and causing societal and economic disruption are alarming. However, the remarkable efforts across the globe pushing for the development and administration of vaccines preventing infection by SARS-CoV-2 is progressing at a rapid pace. In addition to the development of vaccines, research has focused on novel therapeutic strategies, including antivirals, to treat infections. Though significant milestones continue to be achieved concerning the pathogenicity of the novel coronavirus, our knowledge of the molecular mechanisms of its infection, replication, and treatment remains limited.

Coronaviruses are enveloped in a lipid bilayer with a diameter of ~125 nm, which houses the Spike (S) protein along with the membrane (M), envelope (E), and nucleocapsid (N) structural proteins (Lai & Cavanagh, 1997). Much of the current research efforts are focusing on the virus’s pathogenic protein, the S protein, responsible for its human receptor recognition as a target for the development of various therapies and vaccines. Nevertheless, some antiviral therapies are based on simultaneously disrupting several parts of the viral proteome. Thus, the study of other SARS-CoV-2 proteins appears important as well.

Many viruses contain ion channels, generally referred to as viroporins (José Luis Nieva et al., 2012), small hydrophobic proteins that co-assemble to form ion channel pores that permeate ions across the cell membrane or integrate into cellular compartments such as the endoplasmic reticulum (ER) or Golgi apparatus, thereby disrupting different physiological properties of the cell. One of the most widely studied examples of viroporins is the M2 proton channel from the influenza A virus (Pinto et al., 1992), whose structure has revealed detailed insights into the molecular mechanism of the channel, the mechanism of inhibition by anti-influenza drugs, and the effect of mutations causing resistance (Schnell & Chou, 2008; Stouffer et al., 2008). Viroporins are crucial for viral pathogenicity due to their role in different steps of the viral life cycle, including viral replication, budding, and release (Stouffer et al., 2008).

The SARS coronaviruses also contain such viroporins. In the 2003 SARS-CoV, three viroporins, E, ORF3a, and ORF8a were identified, of which E and ORF3a were shown to be required for maximal SARS-CoV replication and virulence (Castaño-rodriguez et al., 2018). The E protein is the smallest (~8.4 kDa) and most enigmatic of the viral structural proteins, and has been reported to form cation-selective ion channels (reviewed in Schoeman and Fielding, 2019; Verdiá-Báguena et al., 2012; Wilson et al., 2004). It was also demonstrated that the E protein contributes to viral pathogenesis and forms a target for antiviral drug development (Jimenez-Guardeño et al., 2014; Nieto-Torres et al., 2014; Wilson et al., 2004, 2006). Characterization of the electrophysiological properties of the E protein have been hampered by the fact that the E protein may not be efficiently targeted to the plasma membrane but to the ER-Golgi compartments (Cohen et al., 2011; Nieto-Torres et al., 2011). Structures have also been elucidated using NMR spectroscopy for peptides corresponding to the transmembrane region or various other truncations, revealing a transmembrane α-helix (Yan Li et al., 2014; Pervushin et al., 2009); suggesting, along with blue-native gels and a very recent solid state NMR structure of the E protein from SARS-CoV-2 (Mandala et al., 2020), that the oligomeric assembly of the E ion channel could be a pentamer, although this is not yet directly validated by structural data for the full-length ion channel.

Almost all studies aiming to circumnavigate the dearth in functional information concerning the E protein have implemented a technique of reconstituting purified proteins or synthesized peptides into artificial lipid bilayers (Castaño-rodriguez et al., 2018; Nieto-Torres et al., 2014; Regla-Nava et al., 2015; Siu et al., 2019). However, since a comprehensive functional characterization of the full-length protein has not yet been performed, the relevance of the E protein rests in its well-documented role as a fundamental pro-inflammatory SARS-CoV virulence factor (Almazán et al., 2013; Netland et al., 2010; Ortego et al., 2002; Regla-Nava et al., 2015). Additionally, several studies have shown that the E protein is essential for viral replication, suggesting that novel inhibitors of the protein could work post-viral entry before new viral particles are able to bud and infect other cells (Castaño-rodriguez et al., 2018).

In the present study, we aimed to achieve a comprehensive biophysical and functional characterization of the E protein from the novel coronavirus. We expressed the E protein from SARS-CoV-2 fused to an EGFP to study the cellular localization of the protein. Combining these with coarse-grained (CG) molecular dynamics (MD) simulations, we also provide evidence for a membrane curvature effect imposed by the E protein, which is compatible with its purported role in budding and virus particle formation. Additionally, our electrophysiological data points towards a pH-sensitive current in HEK293 cells expressing the E protein using the whole-cell patch clamp technique. Furthermore, atomistic MD simulations based on structural models of pentameric E protein resulted in a collapsed state consistent with a non-conducting conformation of the ion channel. Insight into the function and structure of these viroporins provides an interesting avenue to develop therapies with selective modulation against these proteins, namely due to the lack of homology between coronavirus viroporins and human ion channels.

## Materials & Methods

### Cloning and Expression

For microscopy and electrophysiology, the coding sequence for the E protein from SARS-CoV-2 was initially codon optimized for *Xenopus laevis* and synthesized (GenScript, Piscataway, NJ). The gene was later modified for expression in HEK293 cells by adding a fluorescent EGFP tag, in frame, to the C-terminus of the E-protein sequence via overlap extension PCR and subsequently subcloning the product into a pcDNA3.1 vector; generating the EcGFP construct. HEK293T cells were transiently transfected by this construct using Mirus TransIT-293 (Mirus Corporation). Transfected cells were seeded for imaging or electrophysiology experiments 24 hours post-transfection.

### Purification

The full-length E-protein sequence from SARS-CoV-2 (GenBank Accession: NC_045512.2) was purchased from GenScript as a synthetic gene with optimized codon use for expression in *Xenopus laevis*. The gene was subcloned in the pFastBac1 (for Sf9 expression) and pEG-BM (for HEK293 expression) vectors and baculovirus was generated according to the bac-to-bac baculovirus expression system.^85^ Infected cells were harvested 72 hrs post infection and lysed with an EmulsiFlex C5 (Avestin, Ottawa, Canada). Lysate was separated by ultracentrifugation at 100,000 x *g* at 4 °C for 1 hour and resuspended in an equivalent volume of extraction buffer containing 50 mM Tris-HCl (pH 7.5), 200 mM NaCl, 5 mM MgCl_2_, 100 µg/mL DNaseI, 1 mM PMSF, and protease inhibitors (1 µg/mL leupeptin, 1 µg/mL pepstatin, 1 µg/mL aprotinin). Protein was solubilized with the addition of 0.1% (v/v) DDM/CHS (Anatrace) for 2 hours at 4 °C. Solubilized protein was cleared by ultracentrifugation at 30,000 x *g* at 4 °C for 30 min and resuspended in buffer containing 50 mM Tris-HCl (pH 7.5), 200 mM NaCl, 5 mM MgCl_2_, and 0.003% (v/v) DDM/CHS. The solution was then purified by affinity chromatography after incubation for 1 hour on Nickel-Sepharose beads (Cytiva) at 4 °C. The column was washed with 5 column volumes of the same buffer + 40 mM imidazole, and protein was eluted with 3 column volumes of the same buffer + 300 mM imidazole. The eluent was then purified again by affinity chromatography after incubation with anti-GFP beads (GFP-nanobody corresponding to sequence of PDB ID: 3OGO coupled to NHS-activated agarose beads according to manufacturer’s protocol) for 1 hour at 4 °C. The solution was washed with the same buffer (without imidazole). The E protein was cleaved from the EGFP-8xHis fusion by incubation of the beads with 150 units of thrombin (Sigma-Aldrich) overnight at 4 °C. Cleaved E protein was concentrated using a 10 kDa MWCO concentrator (Millipore Sigma) to ~ 1 mL and further purified on a Superdex 200 10/300 Increase (GE Healthcare) column equilibrated with buffer containing 20 mM Tris-HCl (pH 7.5), 150 mM NaCl, 1 mM MgCl_2_, and 0.03% (v/v) LMNG. Peak fractions corresponding to oligomeric fractions were pooled and concentrated to ~2 mg/mL. Samples were collected throughout the purification process, loaded with Laemmli loading buffer, and denatured at 95 °C for 10 min for analysis on a 4-15% SDS-PAGE under reducing conditions.

### Microscopy

HEK293T cells were seeded on an 18 mm diameter coverslip and transiently transfected with the constructs mentioned above (Results and Fig. 2). 24 hr post-transfection, cells were loaded with a dye to visualize the ER (ER-Tracker™ Blue-White DPX; ThermoFisher Scientific) at a final concentration of 0.5 µM for 20 min at 37 °C and 8% CO_2_. Cells were washed with phosphate-buffered saline (PBS), and subsequently labeled with a PM probe (Wheat Germ Agglutinin-Alexa Fluor 633 conjugate (WGA-Alexa633); ThermoFisher Scientific) at a final concentration of 5 µg/mL and incubated for 10 min at 37 °C and 8% CO_2_. Cells were washed with PBS for image acquisition. All images were collected with a Zeiss LSM 880 – Airyscan using a Plan-Apochromat 63x/1.4 Oil DIC M27 objective (Cell and Tissue Imaging Cluster (CIC, KU Leuven), supported by Hercules AKUL/15/37_GOH1816N and FWO G.0929.15 to Pieter Vanden Berghe, KU Leuven. EGFP fluorescence was excited at 488 nm, imaged at 5% laser power and a detector gain of 1100. ER-Tracker fluorescence was excited at 405 nm, imaged at 5% laser power and a detector gain of 553. WGA-Alexa633 fluorescence was excited at 633 nm, imaged at 5% laser power and a detector gain of 870. Colors were added to signals post-acquisition using default look-up tables in Fiji software (Schindelin et al., 2012).

### Calcium imaging

Fura-2-based ratiometric intracellular Ca^2+^ measurements were performed as described previously (Vriens et al., 2011; Wagner et al., 2008). Briefly, cells were loaded with the Ca^2+^ sensitive dye, Fura-2-acetoxymethyl ester (Fura-2AM, Molecular Probes; Invitrogen), in culture medium for 25 min at 37 °C. Experiments were performed in extracellular, Ca^2+^-free KREBS solution containing (in mM): 150 NaCl, 6 KCl, 10 EGTA, 1.5 MgCl_2_, 10 glucose, and 10 HEPES buffered to pH 7.4 (NaOH). Intracellular Ca^2+^ was monitored as the ratio between fluorescence intensities upon illumination at 340 and 380 nm using an MT-10 illumination system and Olympus xcellence pro software (Olympus). After a 5-8-min baseline recording, [Ca^2+^]_ER_ levels were assessed by the addition of 2 µM ionomycin (IO; Calbiochem, San Diego, CA), a Ca^2+^ ionophore. IO-induced Ca^2+^ rises were regarded as [Ca^2+^]_ER_ content that could be estimated from the area under the curve of the [Ca^2+^]_i_ rise.

### Electrophysiology

Whole-cell patch clamp recordings were performed using an EPC-10 amplifier and Patchmaster software (HEKA Elektronik; Lambrecht/Pfalz, Germany). Data were sampled at 5-20 kHz and digitally filtered off-line at 1-5 kHz. Holding potential was 0 mV, and cells were ramped from −150 mV to +150 mV over the course of 800 ms, every 2 s. Pipettes with a final series resistance of 2-4 MΩ were fabricated and filled with intracellular solution. The standard intracellular solution contained (in mM): 130 Cs-Aspartate, 2 Mg-ATP, 10 MgCl_2_, 1 EGTA, and 10 HEPES buffered to pH 7.3 (CsOH). The standard extracellular solution contained (in mM): 135 NaCl, 5 KCl, 1 MgCl_2_, 2 CaCl_2_, 10 glucose, and 10 HEPES buffered to pH 7.4 (NaOH). Recordings to measure the effects of pH used the same standard extracellular solution buffered to pH 6.0, 5.0, and 4.0 with HCl. All measurements were performed at room temperature. Liquid junction potentials were corrected for off-line.

### Coarse-grained simulations

CG simulations were run using GROMACS 2020 (Páll et al., 2020) with a timestep of 20 fs for 20 µs. The standard equilibrium procedure from CHARMM-GUI (Jo et al., 2008) was used before the final production, in which the positions of the protein backbone beads were still restrained for sufficient sampling (except the system with multiple separated monomers). The mean temperature and pressure were kept constant at 310 K and 1 bar using the v-rescale thermostat (tau=1 ps) and the Parrinello-Rahman barostat (tau=12ps) (Bussi et al., 2007; Parrinello & Rahman, 1981). Martini force field version 2.2 for protein and version 2.0 for lipids were used (Marrink et al., 2019).

For the smaller systems with one/no protein. The E-protein-pentamer model (PDB ID: 5×29) was converted to CG model with CHARMM-GUI. It was then embedded into a membrane patch that represents the ER-Golgi lipid composition (59% DOPC, 24% DOPE and 17% DOPS) (Verdiá-Báguena et al., 2012). In total ~2000 lipid molecules were placed inside a rectangular simulation box with the size of 256 Å × 256 Å × 105 Å. The conformation of the E-protein monomer was preserved by applying harmonic restraints with a force constant of 1000 kJmol^−1^nm^−2^ onto the backbone beads of the protein.

For the larger system of E-protein monomers, 20 E-protein monomers (PDB ID: 2MM4) were randomly placed into a simulation box with the size of 512 Å × 512 Å × 105 Å, allowing both sufficient space for protein diffusion and reasonable sampling of protein-protein interaction. In total 240,000 beads were contained in the system. The conformation of the E-protein monomer was restrained with the ELNEDYN elastic network (Periole et al., 2009).

The membrane curvature analyses were performed on the last 10 µs with MemSurfer (Bhatia et al., 2019) tool. The mean local curvature of the smooth approximate surfaces within 25 Å of the proteins for both cytoplasmic and lumenal leaflets were compared and plotted as Raincloud plots (Allen et al., 2019) where each datapoint represents an averaged curvature value for the analyzed frame. Additionally, for the larger system, the mean curvature of the last 400 ns was binned with SciPy (Virtanen et al., 2020) binned_statistic_2d and the last position of monomers were mapped onto the grid. The visualization snapshots were created with VMD (Humphrey et al., 1996).

### Atomistic simulations

Molecular dynamics simulations were run using GROMACS mdrun 2019 (Páll et al., 2020) with a timestep of 2fs, reaching timescales of 300 to 600ns. The mean temperature and pressure was kept constant at 310 K and 1 bar using the v-rescale thermostat (tau=1ps) and the Parrinello-Rahman barostat (tau=5ps) (Bussi et al., 2007; Parrinello & Rahman, 1981). The systems employed the CHARMM36m force field (Jo et al., 2017) or the AMBER99SB-ILDN + Slipids forcefield (Jämbeck & Lyubartsev, 2012, 2013; Maier et al., 2015). Bonds involving hydrogen were constrained using LINCS (Hess, 2008) and the TIP3P (Jorgensen et al., 1983) water molecules were kept rigid with SETTLE (Miyamoto & Kollman, 1992). The van der Waals interactions were switched with “Force-Switch’’ from 10 Å to 12 Å in the case of the CHARMM36m force field, while a simple cut-off of 15Å was used with the AMBER99SB-ILDN + Slipids force field. Some of the simulations were run under a constant 300 mV external electrostatic potential to try to improve stability and water and ion permeation. Long range electrostatics were calculated with the Particle Mesh Ewald method (Essmann et al., 1995). The version of GROMACS used in the simulations was found to contain a bug on the external electrostatic potential feature after running these simulations. We re-ran some simulations with a patched version of GROMACS and found no significant differences. Some of the simulations used the CHARMM-GUI equilibration protocol with an extension of the duration of the steps (Jo et al., 2017). In other cases, pentameric restraints were used (Dämgen & Biggin, 2020). Visualization of trajectories and structures was done with VMD (Humphrey et al., 1996) and data analysis with MDAnalysis (Gowers et al., 2016).

To our knowledge, the only available apparently open structure of a pentameric channel in this family (PDB ID: 5×29 residues 8-65) was determined for the SARS-CoV variant based on solution-NMR constraints and the C40A, C43A and C44A engineered mutations (Surya et al., 2018b). We have restricted our simulation models to the residues for which coordinates are available for every particular PDB entry. Thus, for 5×29 derived models the structure includes the transmembrane domain and part of the C-terminal domain (residues 8-65). For our simulations, this structure was converted to the SARS-CoV-2 wildtype sequence using the SWISS homology model (Waterhouse et al., 2018) of the first model provided in the 5×29 PDB file. Models 6 and 7 of the NMR structure were also used and were converted to the SARS-CoV-2 sequence using PyMOL (Lill & Danielson, 2011). Palmitoylation and N-glycosylation are known post-translational modifications of the E protein (reviewed in Ruch and Machamer, 2012; reviewed in Schoeman and Fielding, 2019). We also performed simulations in which residues C40, C43, C44 were palmitoylated and others where residue N66 was glycosylated with man5 using the previous 5×29-based (residues 8-65) model. Both modifications were added using CHARMM-GUI (Jo et al., 2017) and later manually adjusted to avoid steric clashes. Additional models consisted of the SARS-CoV E-protein monomer structure (PDB ID: 2MM4 residues 8-65) pentamerized using 5×29 as a template, and the more recent solid-state NMR pentameric TM-domain E-protein structure of SARS-CoV-2 (PDB ID: 7K3G residue 8-38) (Mandala et al., 2020). All systems were prepared using CHARMM-GUI (Jo et al., 2017). The proteins were embedded in a 80Å x 80Å POPC, POPS, cholesterol (3:1:1) bilayer using the PPM server and an additional translation of −8Å perpendicular to the membrane plane. Due to the different availability of lipids in CHARMM-GUI for different force fields the AMBER99SB-ILDN + Slipids simulations used POPG lipids instead of POPS. 22.5Å water layers were added on both sides of the membrane with a salt concentration of 75 mM NaCl and 75 mM of KCl.

### Permeation calculations

A fairly stable open conformation was selected from the previous unbiased simulation 8 using the 5×29-based model (Table S1) and submitted for permeation calculations using the accelerated weighted histogram (AWH) method implemented in GROMACS2020 with the CHARMM36m force field. AWH is an extended ensemble method which samples and adaptively optimizes the ensemble, while estimating the free energy (Lindahl et al., 2014).

Two sets of 4 AWH simulations, measuring the permeation of Cl^−^, Na^+^, K^+^ or Ca^2+^, were carried out to estimate the free energy landscape of the passage of each ion through the open pore. The N-terminal unstructured loops (residues 8 to 11) were removed from each monomer to avoid an artificial high energetic barrier (as noticed from a first test of free energy calculations, data not shown) induced by the pore obstruction due to their presence. Models for AWH were built by introducing an ion (Cl^−^, Na^+^, K^+^ or Ca^2+^) within the pore. The whole system was then embedded in a membrane bilayer and immersed in a solvent box with TIP3P water, NaCl and KCl ions using the CHARMM-GUI platform as described in the previous section. Six equilibration steps were performed, before the production AWH runs, by keeping the protein alpha carbons restrained and by gradually decreasing the restraints for the protein heavy atoms and lipids (head groups/dihedral angles).

For each equilibrated system, AWH bias potential was applied on the center of mass z distance between the ion and the residues F26 within the pore center. We used 6 walkers to sample multiple transition pathways within one simulation and thus enhanced the sampling and accelerated the convergence of the simulations. The sampling interval was z ∈ [−6.5, +6.5] and z ∈ [−6.95, +6.95] nm, respectively for the first and the second sets of simulations. A higher pulling distance interval permitted a better convergence of the simulations. To keep the ion close to the pore, the coordinate radial distance was restrained to stay below 1 nm (in the xy plane) from the pore center axis by using a flat-bottom position restraints (cylinder) with a force constant k = 10 000 kJ.mol^−1^.nm^−2^. Alpha carbon position restraints - with force constants k_x_,k_y_,k_z_ = 1 000 kJ.mol^−1^nm^−2^ - were applied to keep the pore channel hydrated and open.

The MD time step was 2 fs. Bonds involving hydrogens were constrained using LINCS. The temperature was kept at 310 K using the v-rescale thermostat and the pressure at 1 bar using Berendsen pressure coupling. Long-range electrostatics were calculated using particle mesh Ewald. Long-range Lennard-Jones interactions were calculated by switching the force to zero for atom distances 10–12 Å. The z direction was set to 0 compressibility. The simulation time was 250 ns and 100/200 ns long for the first and second sets of simulations, respectively. The *gmx awh* module in GROMACS2020 was used to plot the free energy profiles, coordinates and target distributions.

## Results

### Primarily intracellular expression of E protein

We initially performed a sequence alignment to compare the E protein from SARS-CoV of the 2003 and of the novel coronavirus (SARS-CoV-2). While the E protein from SARS-CoV is one residue longer, the two variants remain nearly identical, with three mismatches in the C-terminal domain (Fig. 1A). We also aligned the truncated sequences of the E protein from SARS-CoV and SARS-Cov-2 used for previous NMR-based structure determination, including a truncated peptide encoding residues 8-65 of the SARS-CoV E protein (E Trunc (2MM4/5×29)) (Yan Li et al., 2014; Surya et al., 2018b), and a peptide encoding the transmembrane domain of the SARS-CoV-2 E protein (residues 8-38, E TM (73KG)) (Mandala et al., 2020) (Fig. 1A).

**Figure 1.**
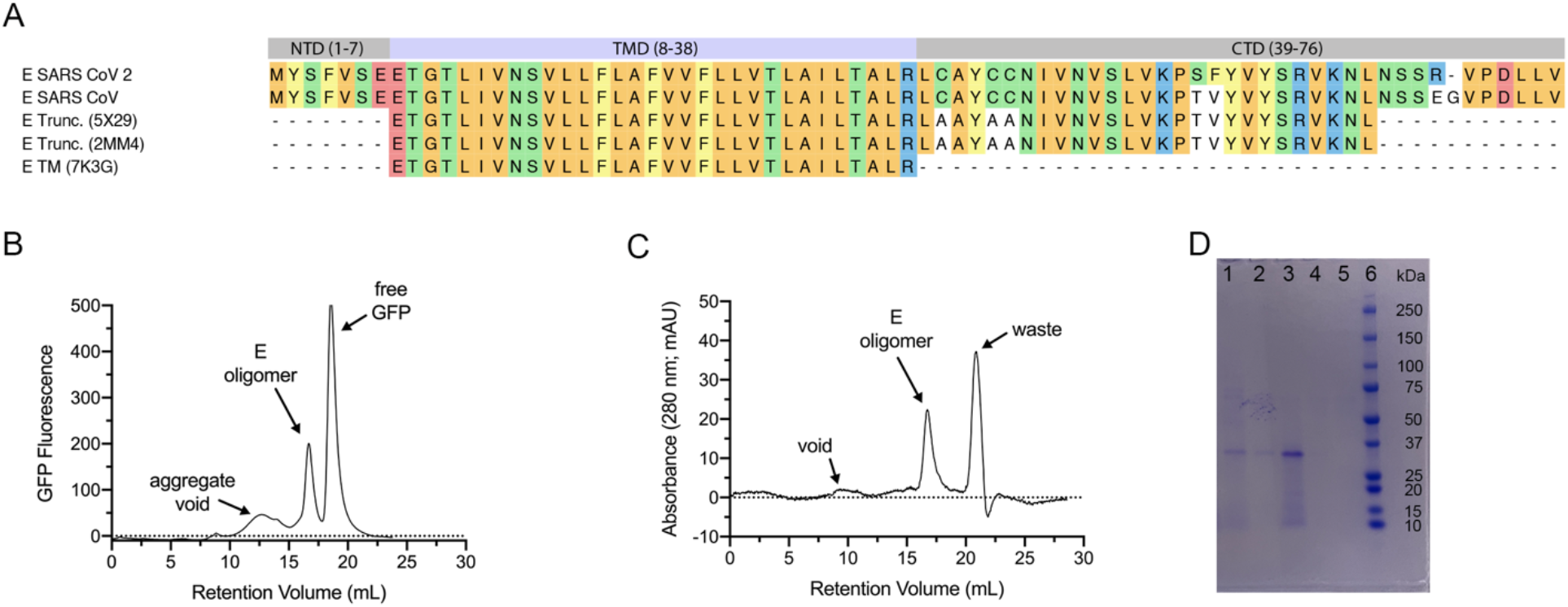
Sequence and oligomerization of SARS-CoV-2 E protein. A: Protein sequence alignment using clustal omega. Sequences (from top to bottom): E protein of SARS-CoV-2; E protein of SARS-CoV; E (SARS-CoV) truncation used for NMR structure (PDB ID: 5×29); E (SARS-CoV) truncation used for NMR structure (PDB ID: 2MM4); E (SARS-CoV-2) transmembrane domain used for NMR structure (PDB ID: 7K3G). Domains of E SARS-CoV-2 are annotated above its sequence, illustrating the amino-terminal domain (NTD, residues 1-7), transmembrane domain (TMD, residues 8-38), and carboxy-terminal domain (CTD, residues 39-75). Colored residues match E SARS-CoV-2 reference sequence; mismatches are not colored. B: Chromatogram result from FSEC on Superose 6 10/300 column where y-axis represents GFP fluorescence and x-axis represents retention volume (mL). Corresponding fractions are estimated by arrows. C: Chromatogram result from SEC on Superdex 200 10/300 Increase where y-axis represents UV absorbance (280 nm) and x-axis represents retention volume (mL). Corresponding fractions are estimated by arrows. D: SDS-PAGE loaded with coomassie-stained samples from SEC purification under reducing conditions. Lane 1: SEC load (diluted 1:2); 2: E oligomer fraction (V_R_ = 16.8 mL); 3: concentrated E oligomer fraction; 4: waste fraction (V_R_ = 20.9 mL); 5: concentrated waste fraction; 6: marker with corresponding mass denoted to the right of the band (kDa).

Here, we report on the full-length E protein from SARS-CoV-2. We synthesized this gene and incorporated a cleavable EGFP and 8x-histidine affinity tag for recombinant expression and purification in Sf9 cells. After expression, we solubilized the E protein with a 10:1 mixture of n-dodecyl-β-D-maltoside and cholesteryl hemisuccinate (DDM/CHS), and using fluorescence-based size exclusion chromatography (Kawate & Gouaux, 2006) equipped with a Superose 6 column, we observed that the E protein expresses and forms stable complexes (Fig. 1B). After cleaving the affinity tags, we loaded the purified E protein onto a size exclusion column equipped with a Superdex 200 10/300 Increase column, and observed that the protein elutes as a monodisperse peak around ~16.5 mL (Fig. 1C), which corresponds to an oligomeric band of ~35 kDa on SDS-PAGE (Fig. 1D). Our results here show for the first time, to our knowledge, that the full-length E protein from SARS-CoV-2 is capable of multimeric assembly independent of ligands or other factors.

Next, we synthesized a covalently-linked EGFP gene to the C-terminus of the E protein to visualize where the novel viroporin was being trafficked or compartmentalized in HEK293T cells. We transfected this construct and imaged 24 hours post transfection using confocal microscopy (Fig. 2A). Our results were consistent with previous reports examining the localization of the E protein from other CoVs, showing that the majority of the E protein remains intracellular, most likely around the ER and ER-Golgi intermediate compartment (ERGIC), as shown by the protein’s colocalization with the ER-Tracker™ signal (Fig. 2A).

**Figure 2.**
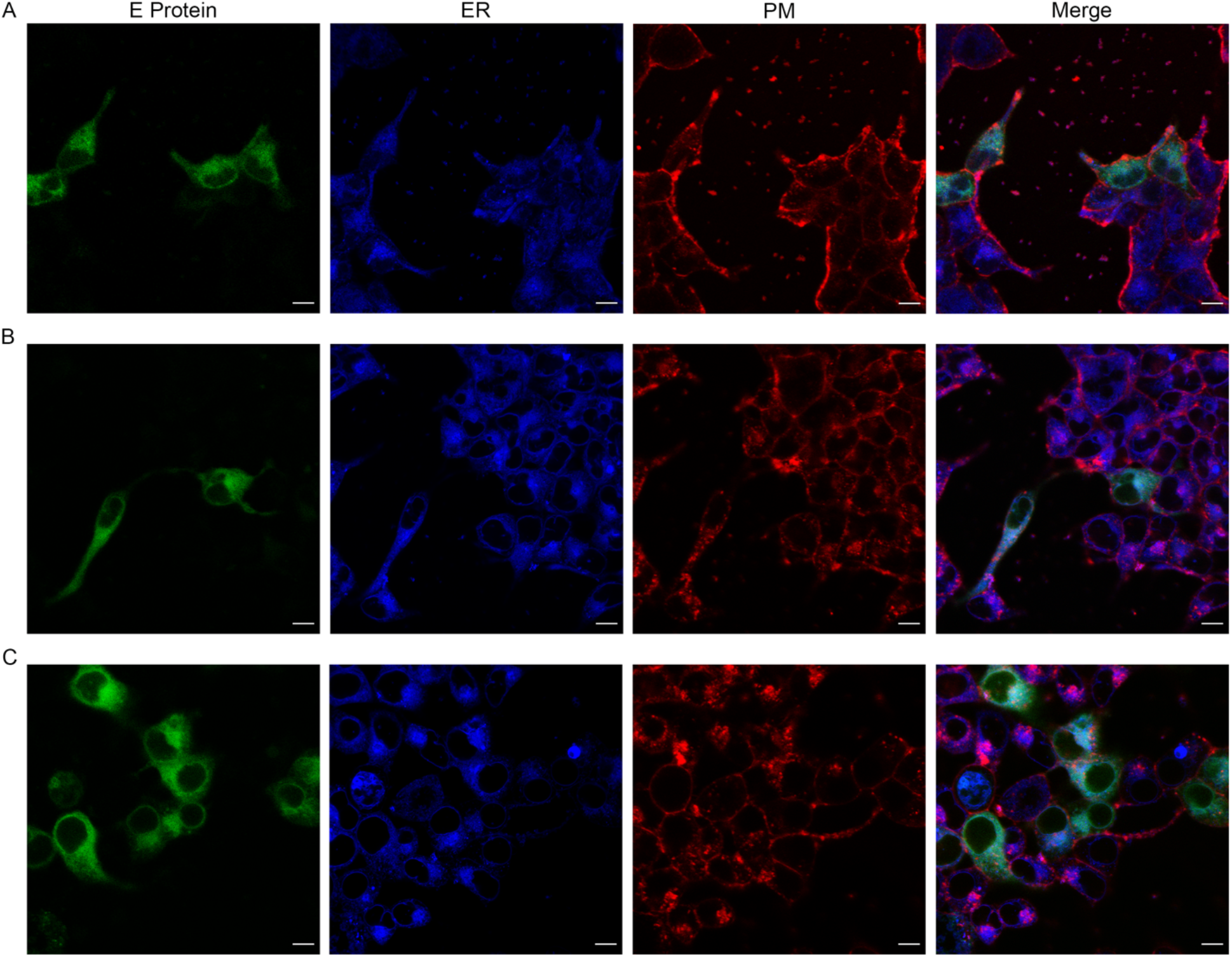
E protein localizes intracellularly in HEK293T cells. Live-cell confocal microscopy shows the E protein localizing to intracellular compartments. A: E protein with a C-terminal EGFP fusion. B: E protein with a N-terminal export signal sequence from the α7 subunit of nAChR and C-terminal EGFP fusion. C: E protein containing point mutation P54A with a C-terminal EGFP fusion. HEK293T cells were transiently transfected with the respective constructs and imaged 24 hpt (1^st^ column, E protein, green signal). Cells were also loaded with an ER-Tracker™ to visualize the ER compartment (2^nd^ column, ER, blue signal). Cells were also loaded with PM probe (see Methods) (3^rd^ column, PM, red signal) to visualize the PM. 4^th^ column represents merged signals. Scale bars represent 10 µm.

It is well-known that many proteins encode export or retention signals, including the α subunits of the nicotinic acetylcholine receptor, which houses a signal peptide on its N-terminal extracellular domain that aids in translocating the protein to the plasma membrane (Dash et al., 2014). To enable us to better perform functional studies of the E protein, we attempted to improve the protein’s expression at the plasma membrane (PM) by synthesizing a modified gene with this signal peptide from the α7nAChR subunit incorporated at the N-terminus of the E protein to examine if there would be an increase in surface expression; however, we did not observe any marked improvement in plasma membrane expression (Fig. 2B). Additionally, previous reports studying the localization of envelope proteins from various CoVs indicate conserved retention motifs; namely a beta-proline-beta motif in the C-terminus that acts as an ER retention signal (Cohen et al., 2011; Yan Li et al., 2014). We performed single point mutagenesis to switch this conserved proline at position 54 to an alanine in an attempt to eliminate the retention signal. However, in contrast to previous reports expressing this mutated construct in the E protein from SARS-CoV, or inserting more extensive Golgi-export signals from mammalian channels (Cabrera-Garcia et al., 2021), we did not observe any significant improvement in PM expression for the novel protein (Fig. 2C). Pearson’s coefficients revealed slightly higher, but not statistically significantly different, correlations between E-Protein and the ER, than between E-Protein and the PM (*r*_E:ER_ = 0.32 ± 0.13 vs. *r*_E:PM_ = 0.22 ± 0.21), even with modification to the signal sequences (Fig. 2B-C, Fig. S2A-B). These results suggest that the limited surface expression may be consistent with the purported role of the E protein in virus budding at the ERGIC.

### E protein induces membrane curvature in coarse-grained simulations

Budding is an important stage of the viral life cycle (José Luis Nieva et al., 2012), and the E protein is hypothesized to induce bending of the membrane, which would play a pivotal role in the process (reviewed in Schoeman and Fielding, 2019). To investigate this, we carried out coarse-grained MD simulations, enabling us to model membrane systems large enough to be able to undergo structural deformation over the multi-microsecond simulation length. We then measured the local membrane curvature around an E-protein pentamer (Fig. 3A) derived from an NMR structure of a portion of the SARS-CoV E protein (PDB ID: 5×29) (Surya et al., 2018b). The E-protein pentamer significantly bent the membrane for both cytoplasmic and lumenal leaflets in a fashion that could facilitate viral budding, compared to a pure lipid-bilayer patch, the E-protein monomer, or the ORF3a protein (Fig. 3B-C).

**Figure 3.**
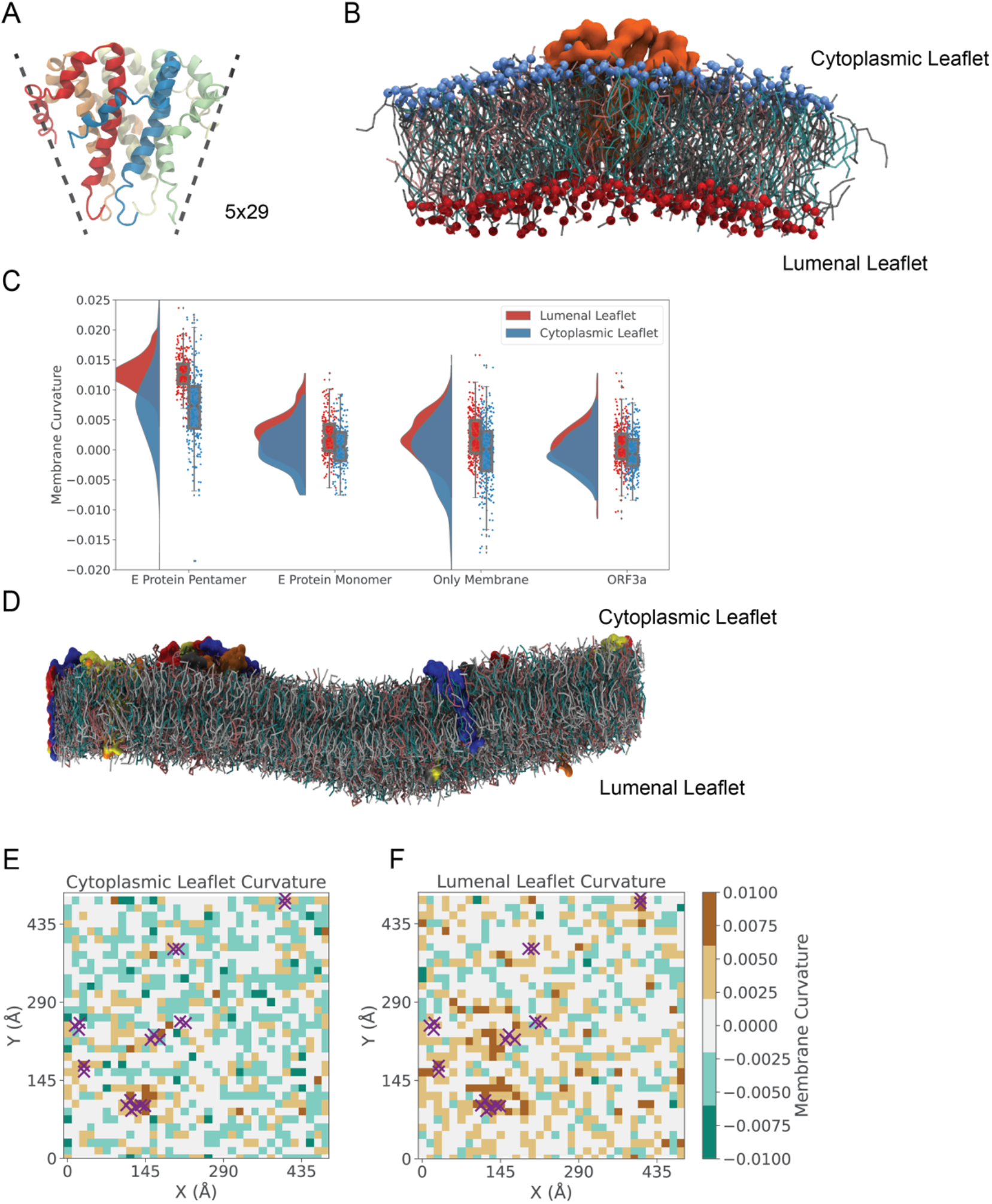
E protein induces membrane curvature in coarse-grained simulations. A: Model of the E protein, represented by the cone-shaped pentameric solution-phase NMR structure covering residues E8–L65 of the SARS-CoV variant (PDB ID: 5×29). B: Final snapshot of the CG system with the E-protein pentamer embedded in a mixed lipid bilayer. For clarity, only lipids within 50 Å from the protein were displayed. C: The mean local membrane curvature within 25 Å of the protein for both cytoplasmic and lumenal leaflets, with probability distribution on the left, and raw data (N = 500 samples of trajectory frames) with box plots indicating the median, interquartile range (25^th^–75^th^ percentiles) and minimum–maximum ranges on the right. The E-protein pentamer induced a positive bending of the membrane around it while the membrane curvature of the E-protein monomer, ORF3a protein, or pure membrane patch remained planar. D: Final snapshot of the system with 20 E-protein monomers. While the monomers did not form a symmetric pentamer, the major aggregated cluster of monomers did bend the local membrane. Proteins are shown in surface representation, with lipids as sticks (gray: DOPC, cyan: DOPE, pink: DOPS) E-F: Cytoplasmic- and lumenal-leaflet curvature of the last snapshot of the system containing 20 E-protein monomers. The membrane around the major protein cluster displayed the highest local curvature. The positions of protein monomers are indicated by purple crosses.

To test the role of the cone shape of the initial E-protein model in driving membrane curvature, we performed a simulation with a bilayer of the same composition, inserting instead 20 parallel E-protein monomers (PDB ID: 2MM4) randomly in the membrane. Although symmetric pentamers did not form over the 20 µs simulation time, the aggregation of monomers induced bending of the membrane around the major cluster (Fig. 3D-F). This suggests that the cone-shaped structure of the pentameric assembly is not actually required for membrane bending, and that the topology of the monomer featuring a transmembrane and interfacial amphiphilic helix could instead be an important determinant of membrane bending. While the precise budding mechanism of the E protein from SARS-CoV-2 remains elusive, these CG models provide an initial observation of the important role of the protein on membrane bending.

### Intracellular calcium depletion in E-protein transfected cells

Animal viruses are adept at tailoring the cell’s innate Ca^2+^ toolkit to provide sufficient opportunities for the host cell to adjust to the virus infection (reviewed in Zhou et al., 2009). In general, virus infection elicits an increase of intracellular Ca^2+^ levels as a result of altered plasma membrane permeability as well as changes in membrane permeability of internal Ca^2+^ stores (Agirre et al., 2002). To evaluate whether this phenomenon was present in E-transfected cells, we used a Ca^2+^ ionophore, Ionomycin (IO), to empty all Ca^2+^ stores in cells transfected with the E protein (Fig. 4A-B). We observed that the total Ca^2+^ content, determined by the area under the curve (AUC), was decreased by about 61.5% in cells transfected with the E protein (0.1286 ± 0.0745 AU, N = 22) compared to non-transfected cells (0.2002 ± 0.096, N = 19; p = 0.01) (Fig. 4C). The amplitude was also significantly diminished in E-transfected cells (E: 0.04649 ± 0.0338 vs. NT: 0.1036 ± 0.0582, p < 0.001) (Fig. 4C). These results illustrate that cells transfected with the E protein show a depletion of Ca^2+^ upon IO-induced release of intracellular Ca^2+^ stores, suggesting the protein plays a role in leaking, suppressing, or sequestering Ca^2+^ from multiple standard compartments.

**Figure 4.**
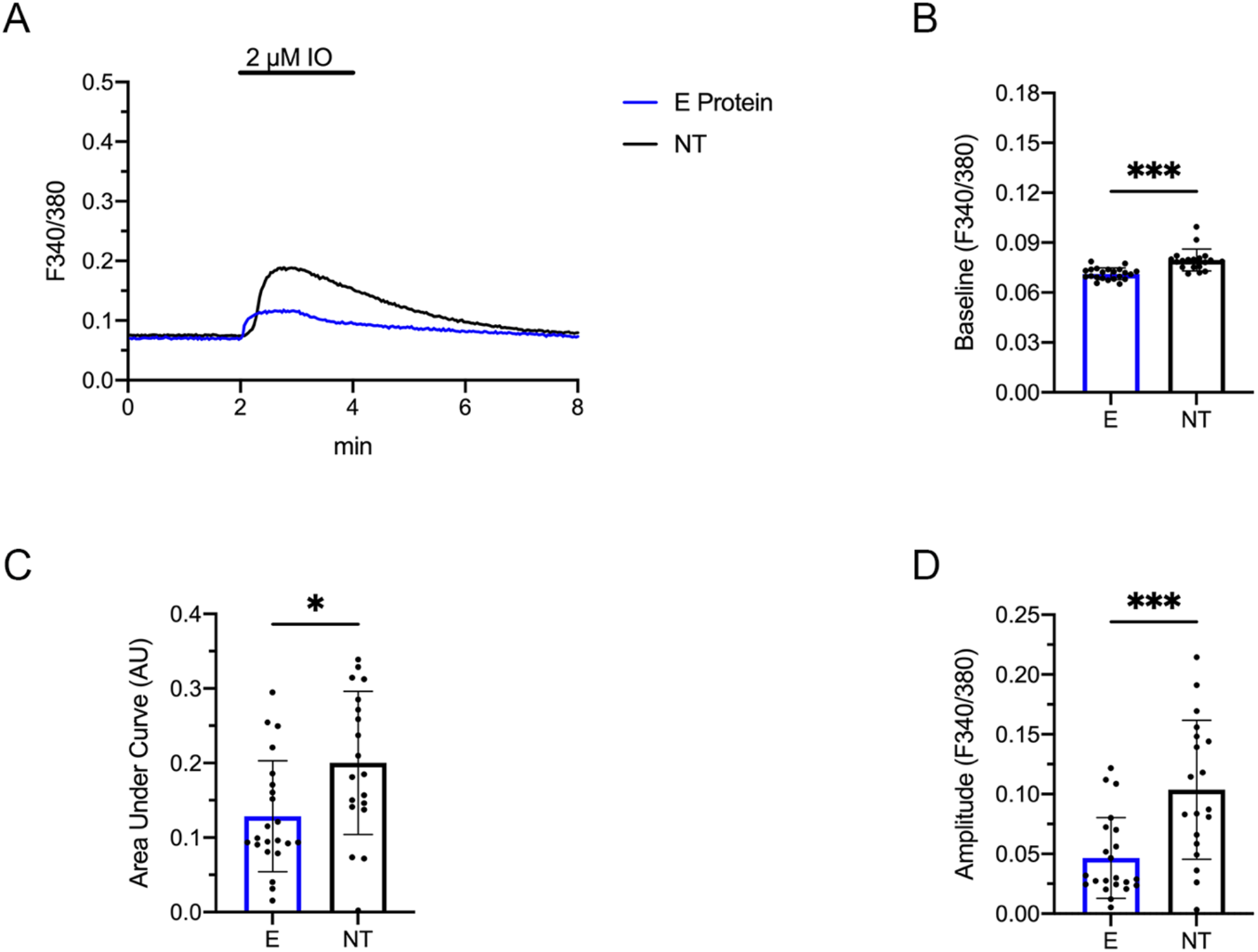
Ca^2+^-store content is diminished after transfection with E protein. A-B: Representative [Ca^2+^]_ER_ measurements in HEK293T cells transfected with the EcGFP construct (A, blue trace) and non-transfected (NT) cells (A, black trace). Recordings were performed in Ca^2+^-free media. 2 µM ionomycin (IO) added after 2 minutes. B-D: Quantification of parameters to evaluate Ca^2+^ store content release. E protein: N = 22; NT: N = 19. B: Baseline, p < 0.001(***). C Area under Curve (AUC), p = 0.01(*). D: Amplitude, p < 0.001(***). Statistical significance was calculated using Welch’s unpaired t-test to account for unequal standard deviations.

We then looked to further characterize the function of the protein as a viroporin. Many human viruses contain pore-forming proteins to modulate cellular functions and regulate viral functions at different stages of its life cycle. While they tend to be small proteins of ~60-120 amino acids in length, their absence does attenuate viral fitness and its pathogenic effects (DeDiego et al., 2007; Wang et al., 2012; Watanabe et al., 2009). Viroporins from other viruses have been shown to transport mainly cations such as protons (H^+^), sodium (Na^+^), potassium (K^+^), and calcium (Ca^2+^) (Lu et al., 2006; Mould et al., 2003; Pham et al., 2017). To examine the permeability of these cations in the E protein, we patch clamped HEK293T cells transfected with the E protein, and applied a voltage ramp from −150 mV to +150 mV. We observed no change in current in response to extracellular buffers containing Na^+^, K^+^, or Ca^2+^ (data not shown). We observed a similar response from two-electrode voltage clamp recordings in *Xenopus laevis* oocytes microinjected with E-protein cRNA (data not shown). Recent studies, on the other hand, have reported a pH-dependent current-voltage relationship where outward currents are elicited in acidic buffer conditions (Breitinger et al., 2021; Cabrera-Garcia et al., 2021). Given the lack of current response to mono- or divalent cations, we investigated the channel’s sensitivity and/or permeability to protons by decreasing the pH of our standard extracellular buffer and measuring the current in response to a voltage ramp. From a pH of 6 and lower, we saw robust, outwardly rectifying currents, reaching a current density at 100 mV of 86.9 ± 48.9 pA/pF in buffer equilibrated to pH 4 (N = 19) (Fig. S1A). Though these current densities appeared to be higher in E-transfected cells compared to non-transfected cells (52.2 ± 58.5 pA/pF; N = 10), the difference was not statistically significant (p = 0.08) (Fig. S1B-C).

Since HEK cells harbor an endogenous, pH-sensitive anion channel, TMEM206 (Lambert & Oberwinkler, 2005), we further characterized the current we observed in E-transfected cells to confirm its identity. In response to voltage steps from −160 mV to +40 mV, we observed outward currents in buffer conditions at pH 4 with similar kinetics to the endogenous TMEM206 (Fig. S1D-E) (Ullrich et al., 2019). A time course of the current response after changing the extracellular buffer pH from 7.4 to 4 showed a fast increase in outward current and a relatively quick decay. These results were also similar to that of TMEM206 in similar conditions (SI Fig. 1F). Ullrich et al., also identified TMEM206 currents could be blocked by pregnenolone sulfate (PS), an endogenous neurosteroid. We thus tested the effect of 50 µM PS in pH 4 conditions in E-transfected cells and non-transfected cells, and observed that PS could effectively reduce the outward current by 88% in E-transfected cells and 95% in non-transfected cells (SI Fig. 1G). Although we cannot rule out a proton response of E-protein channels, we hypothesize that the marginally higher current in E-transfected cells could be due to an off-target upregulation of endogenous channels post-transfection. These observations emphasizing the importance of caution in interpreting pH-sensitive currents in transfected cells.

### Limited stability and permeation in NMR-based models

Given that our reconstitution, expression, CG, and calcium-transport experiments supported a functional role for the E protein in intracellular regulation, we finally used atomistic MD simulations to probe models of the E-protein structure embedded in model lipid bilayers and their dynamics. MD simulations require an initial structure of the studied protein obtained from experiment or homology modeling, and in many cases, the results of the simulations are predicated on the quality of the starting structure. At the time of this work, we identified three plausible templates for the SARS-CoV-2 E protein: a pentameric solid-state NMR ensemble representing the transmembrane domain in an apparent closed state (residues E8–R38, PDB ID: 7K3G), and monomeric and pentameric ensembles derived from solution-NMR studies of the SARS-CoV homolog (residues E8–L65, PDB IDs: 2MM4 and 5×29, respectively). These structures lack the PDZ binding domain at the C-terminal domain (residues 70-75) for which recent structures of this peptide bound to its cognate receptor PALS1 PDZ domain were recently released (PDBs: 7M4R and 7NTK) (Chai et al., 2021b; Javorsky et al., 2021). Although this domain could have potential implications in ion-permeation and/or membrane curvature, we consider this to be an unlikely scenario. Indeed, the E protein is expected to mediate host immune responses through two distinct mechanisms: the transmembrane domain is associated with the activation of NLRP3 inflammasome (Nieto-Torres et al., 2015), while the C-terminal domain is related to PDZ-binding function (Jimenez-Guardeño et al., 2014; Teoh et al., 2010). We thus opted to exclude these residues from our protein models.

The solid-state NMR SARS-CoV-2 E protein (PDB ID: 7K3G), representing only the transmembrane helices with a 3_10_-helix transition at the N-terminal end (residues F20–F23) (Mandala et al., 2020) was simulated using the CHAMM36m and the AMBER99SB-ILDN + Slipids force fields, to characterize the effect of the force fields (see Table S1), both in the presence and absence of a 300-mV potential on the C-terminal side. The resulting set of combinations of protein models, starting conditions, force fields and membrane potentials serves both to discard a potential biasing of the simulations by our modelling choices and to find conditions where conduction is observed. The principal model was equilibrated with pentameric pore restraints as previously reported for other pentameric ion channels (Dämgen & Biggin, 2020). Throughout all simulations, the pore was dehydrated, consistent with a closed or nonfunctional state of the channel. All simulations deviated dramatically from the starting model (4–7 Å RMSD, Fig. S3A) and lost their initially pentameric symmetry. Indeed, in all cases the protein tilted substantially to form an acute angle with the membrane normal, and twisted with respect to the symmetry axis (Fig. S3). Our simulations also showed that the 3_10_ helical twist relaxed to a more classical alpha-helix (Fig. S3B), even in the AMBER99SB-ILDN force field, which has been reported to stabilize 3_10_ helices (Patapati & Glykos, 2011).

The solution-NMR SARS-CoV E protein (PDB ID: 5×29) represented a larger portion of the sequence with more classical alpha-helical transmembrane helices, and with a pore possibly wide enough for ion conduction, providing a plausible model to investigate permeation as well as stability. The homology between SARS-CoV and SARS-CoV-2 allowed us to obtain suitable homology models for SARS-CoV-2 using 5×29 as a template (Methods). We therefore explored this system using several ensemble members, equilibration protocols, force fields (CHARMM36m and AMBER99SB-ILDN + SLipids), and post-translational modifications (Table S1). With only one exception, a common pattern emerged, where after the release of the restraints, the hydrophobic pore of the E protein dehydrated and the protein structure rapidly lost its initial symmetry (4–7 Å RMSDs). Simulations based on a pentameric assembly of the monomeric SARS-CoV E protein (PDB ID: 2MM4) also showed poor structural stability, in addition to pore dehydration (Table S1). The most stable model was obtained from a SWISS-model homology model of the first ensemble member of 5×29 (residues 8-65) in the CHARMM36m force field, with no post-translational modifications; among three simulations carried out under these conditions, one trajectory using pentameric equilibration restraints (Dämgen & Biggin, 2020) and a 300 mV transmembrane potential retained a relatively stable, hydrated pore (Fig. 5A-B). Several computational electrophysiology simulations starting from this model eventually dehydrated; therefore, we can conclude that our most stable simulation only retains a conductive pore on the hundreds-of-nanoseconds timescale. Although it is possible to envision the dehydrated channel undergoing a spontaneous conformational change to a conductive state, the timescale of this type of rearrangements is far greater than those reached by most molecular dynamic simulations.

**Figure 5.**
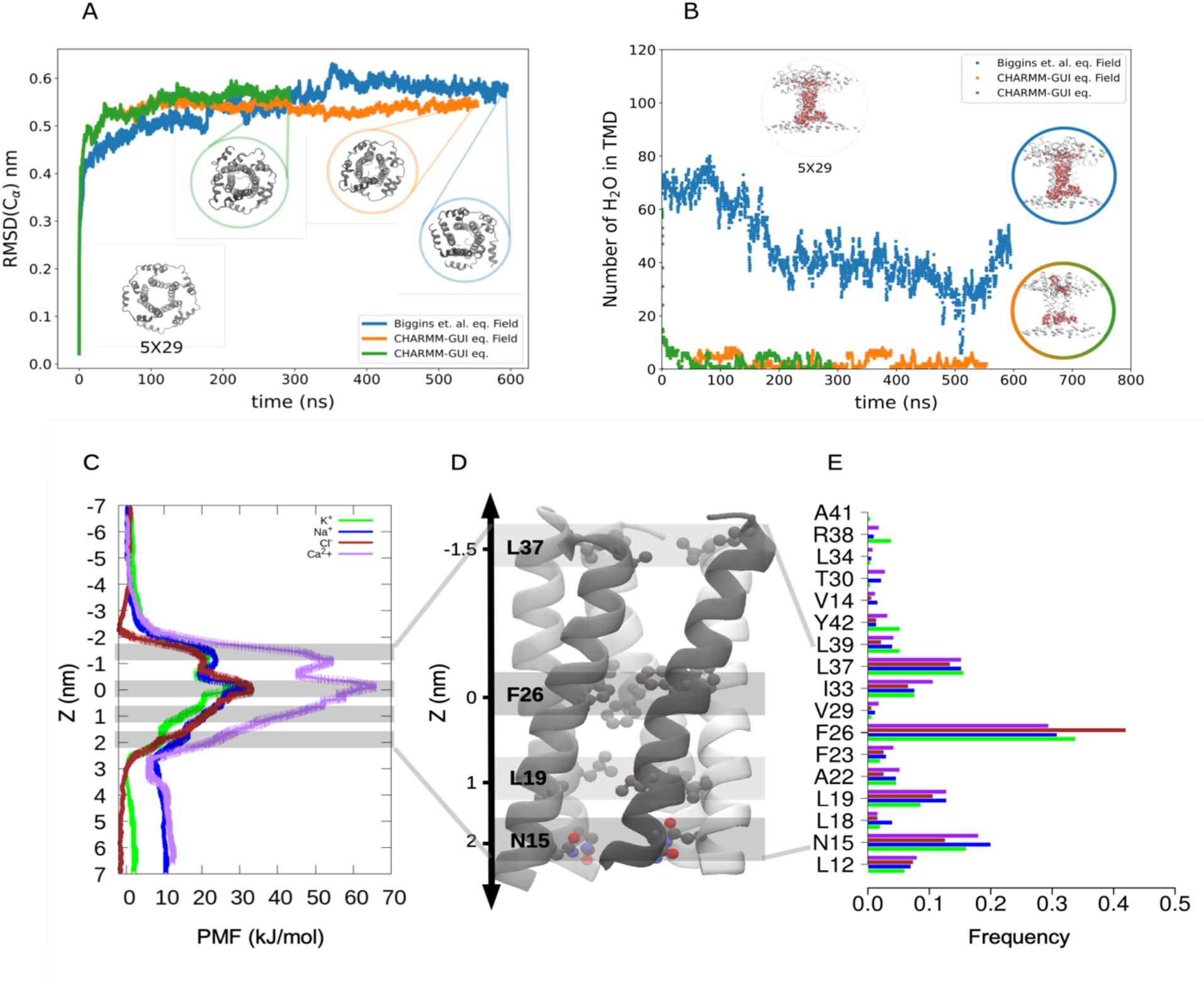
Limited stability and permeation in NMR-based models. A: Root mean-squared deviations (RMSDs) of Cɑ atoms as a function of time during the three most stable simulations of E-protein models based on the pentameric solution-NMR structure (PDB ID: 5×29). Models were equilibrated using the standard CHARMM-GUI protocol and simulated in the absence (green) or presence (orange) of an electric field, or using a pentagonal pore-restrained protocol^38^ and simulated with a field (blue). Inset snapshot at lower left shows the starting model, viewed from the C-terminal side perpendicular to the membrane; additional insets show snapshots from the end of the three simulations. The membrane and solution are omitted for clarity. B: Number of water molecules in the TMD as a function of time during the three simulations shown in *A*. Inset at top left shows the starting model, viewed from the membrane plane; additional insets show representative snapshots from the end of each simulation. The membrane and solution are omitted for clarity. C: Free-energy profiles (mean ± standard deviation) for permeant ions (Cl^−^, orange; Na^+^, blue; K^+^, green; Ca^2+^, purple) along the channel axis of the most stable simulation endpoint in *A–B*. The Z-axis is centered with the highest barrier, proximal to F26 at the channel midpoint, at 0 nm. D: 3D structure highlighting residues in relatively high contact with permeant ions (N15, L19, F26, L37). E: Histogram showing contact frequencies for transmembrane-helix residues in contact with Cl^−^, Na^+^, K^+^ and Ca^2+^ in orange, cyan, green and purple, respectively.

To further probe the ability of the most stable E-protein model (Table S1) to conduct ions, we carried out accelerated-weight histogram (AWH) simulations to assess the permeation and relative selectivity of four ions, including chloride (Cl^−^), K^+^, Na^+^ and Ca^2+^. Starting from the final coordinates of the most stable 5×29 atomistic MD simulation, subsequent pore collapse was prevented by Cα harmonic restraints, and permeating ions were allowed to explore the conformational landscape in a 10-Å radius cylinder around the central pore axis. Over the course of the simulations, the pore remained hydrated and several ion passages were observed (Fig. S4). The one-dimensional free-energy profiles, calculated using adaptive biasing along the Z-axis, showed a principal barrier at the midpoint of the channel axis (Fig. 5C). This position corresponded to residue F26, the most frequent contact (≥30%) for all ions; other contacts (≥10%) included N15 at the N-terminal entrance to the pore and L37 at the C-terminal entrance (Fig. 5D–E). The principal F26 barrier exceeded 20 kJ/mol for both monovalent and divalent ions (Fig. 5C); the free-energy barrier for Ca^2+^ permeation was 67 kJ/mol, higher even than monovalent ions and incompatible with conduction. Thus, stability and permeation profiles of available E-protein models showed little correspondence with functional profiles from coarse-grained simulations and calcium imaging, suggesting a need for further structural characterization.

## Discussion

### Localization in human cells & implications for virus assembly

The findings described above shed light on functional and structural properties of the SARS-CoV-2 E protein using a range of biophysical methods, emphasizing potential pathways for future investigation or development, while also highlighting challenges or gaps in existing knowledge and structure-function models. Our first goal was to confirm the localization of our E-protein GFP fusion construct in transfected HEK293T cells. Although previous reports studying envelope proteins of various viruses have pointed to intracellular localization, this property could not be assumed for coronaviruses. CoVs are distinct in that they bud into the ERGIC prior to export through the host secretory pathway (Hogue & Machamer, 2014). From our confocal data, we observed that the protein was indeed intracellular even in the context of biochemical fusion partners, and overlapped with the fluorescent signal recognizing the ER. This result confirms not only that the E protein from the novel coronavirus may serve similar biological roles as its predecessors, but that our fluorescently tagged fusion protein does not disrupt intracellular expression or localization.

Also consistent with a strong intracellular preference, targeting the E protein to the PM for comprehensive electrophysiological characterization proved challenging, as the export-signal additions and retention-signal mutations implemented in this work did not produce definitive evidence of surface function. It does remain possible that there are other retention motifs present in the sequence such as the DLLV sequence at the C-terminus, as was evidenced recently (Chai et al., 2021a; Pearson et al., 2021; Zhu et al., 2021); however, since this purported retention signal is part of the PDZ domain involved in protein-protein interactions, we did not pursue this signaling motif. Precise identification of the signaling elements responsible for E-protein targeting would provide insight into how CoVs interact with and take advantage of host machinery to assemble new virions, which merits further investigation. In fact, promising results in this regard were reported by another group during preparation of this manuscript (Cabrera-Garcia et al., 2021).

### Oligomerization of purified SARS-CoV-2 E protein

Despite the preferential localization documented above, overexpression and purification of a GFP/octahistidine-tagged E-protein construct in Sf9 insect cells enabled isolation of a monodisperse oligomeric complex. These results complement earlier biochemical reports that the isolated SARS-CoV variant forms multimers, possibly pentamers (Surya et al., 2018a; Torres et al., 2006; Wilson et al., 2006). Although the precise stoichiometry of detergent-solubilized E protein cannot be determined definitively by gel filtration, analysis of our peak SEC fraction coincides with previous reports of the E Protein forming higher order oligomers.

Notably, NMR studies presuming pentameric assembly of SARS-CoV or SARS-CoV-2 E proteins have produced divergent structures. In dodecylphosphocholine (DPC) micelles, the SARS-CoV variant was reported as a pentameric assembly with helical left-handedness, where the side chains of residues V25 and V28 could act as a channel gate (Pervushin et al., 2009). A more recent study, in lyso-myristoyl phosphatidylglycerol (LMPG) micelles, reported an interhelical orientation for the side chain of V25 and helical kink with an overall right-handedness (Surya et al., 2018a). The E-protein transmembrane domain also contains three sequential phenylalanine residues, spaced three residues apart from each other. Surya et al. reported a protein state in which the aromatic side chains of these residues are positioned towards the lumen of the pore. In comparison, the most recently published solid-state NMR spectroscopy-derived structure of the E protein from SARS-CoV-2 reveals a continuous state with these side chains oriented away from the pore and a closed state where the middle aromatic residue, F26, rotates inward, thereby constricting the pore (Mandala et al., 2020). Such widely varying properties of reported structures highlight the need for further biophysical characterization of full-length E-protein assembly, structure, and membrane-transport function, if any.

### Coarse-grained simulations model E-protein effects in a realistic cell membrane

Our CG simulations illustrated the capacity of E protein to induce a degree of membrane curvature, consistent with a role in membrane budding during viral replication and assembly. With few exceptions, lipid diffusion around membrane proteins is too slow to sample efficiently on the timescales of atomistic MD simulations (Wassenaar et al., 2014); accordingly, CG simulations have been applied to study realistic cell-membrane dynamics (Marrink et al., 2019). These methods can accurately measure or predict binding of specific lipids to proteins (Corey et al., 2019) and can, on a larger scale, model changes in shape of biological membranes (Pezeshkian & Marrink, 2021). We captured the membrane-curvature dynamics induced by the E protein by embedding it into a bilayer broadly mimicking the composition and charge of the Golgi membrane. In CG simulations, the protein’s secondary structure needs to be maintained by restraining either the backbone interactions, or the overall conformation of the protein using elastic network approaches (Periole et al., 2009); such restraints limit the ability of CG simulations to probe the dynamics of the protein itself. It should also be noted that protein-protein interactions are exaggerated in the Martini 2 force field (Javanainen et al., 2017), which could explain why disordered clusters of monomers irreversibly formed instead of symmetric pore-like assemblies. Furthermore, although we used a lipid mixture that mimics a realistic membrane composition, other viral proteins were absent, such that we could not capture possible interactions with, for example, the M or S proteins (Boson et al., 2021; Monje-Galvan & Voth, 2021).

Owing to its name, mature coronavirus particles take on a spherical shape, due in considerable part to the assembly of the virion envelope at the ERGIC (Hogue & Machamer, 2014; Westerbeck & Machamer, 2015). During assembly, CoV M and E proteins contribute to producing and pinching off virus-like particles (VLPs) (Baudoux et al., 1998; de Haan et al., 2000; Klumperman et al., 1994). This phenomenon was further confirmed in a recent study illustrating that expression of both M and E proteins also regulates the maturation of N-glycosylation of the S protein (Boson et al., 2021). Since this event usually occurs in the Golgi, it remains possible that the presence of M and E proteins could alter the function of glycosyltransferases (Rosnoblet et al., 2013). In fact, in the absence of the E protein, recombinant CoVs deviate from their typical morphologies, producing propagation-deficient virions (Boson et al., 2021; DeDiego et al., 2007; Lim & Liu, 2001; Ortego et al., 2002). Nevertheless, since CoVs are still capable of assembling without the E protein, the direct role of the E protein in the broader setting of virus infection points more towards inducing a favorable membrane environment into which viral prodigy can insert. Notably, envelope proteins have recently been shown to slow the secretory pathway (Boson et al., 2021; Denolly et al., 2017). Another possibility rises in respect to the E protein’s C-terminal domain, which is reported to harbor a propensity for inducing amyloidogenesis (Ghosh et al., 2015; Mukherjee et al., 2020). Previous studies concerning amyloid proteins of high molecular weight conformers induce a membrane-associated change in local curvature (Burke et al., 2013). However, since the exact topology of the E protein remains unclear, we cannot presently deduce the precise mechanism by which this event occurs, although interesting speculations have been proposed (Duart et al., 2020, 2021; reviewed in Schoeman & Fielding, 2019).

### Implications for calcium homeostasis and viral porin function

The dynamic between a virus and the host cell’s Ca^2+^-signaling pathways and other Ca^2+^-dependent processes is evidenced by direct and/or indirect imbalances in Ca^2+^ homeostasis parameters resulting from affected membrane permeabilities, sequestration of Ca^2+^, and/or Ca^2+^-regulated virus-host interactions. Elevated cytosolic Ca^2+^ may benefit the virus by prompting mitochondrial uptake and thus elevating ATP production to meet higher demands for viral replication (Yanchun Li et al., 2007; Sharon-Friling et al., 2006). Concurrently, an acceleration of Ca^2+^-dependent enzymatic processes may induce Ca^2+^-dependent transcription factors to promote virus replication, as is seen with HIV-1, HCV, and HTLV-1 among others (Bergqvist & Rice, 2001; Ding et al., 2002; Kinoshita et al., 1997). Similarly, a decrease in Ca^2+^-store content could drive inhibition of protein trafficking pathways, hampering the innate immune responses and allowing the virus to escape premature clearance by the host (Doedens and Kirkegaard, 1995; Van Kuppeveld et al., 2005; reviewed in Zhou et al., 2009). A potentially instructive example is the well-characterized nonstructured protein from the enterovirus family, which causes a decrease in ER Ca^2+^ by assembling into pore-forming units, permeabilizing the membrane, and eliciting Ca^2+^ efflux (Irurzun et al., 1995; José L. Nieva et al., 2003).

Although a previous study reported the E protein from SARS-CoV to permeate Ca^2+^ (Nieto-Torres et al., 2015), we furthered these findings and observed that transfection with E protein lowered cytosolic Ca^2+^ compared to non-transfected cells. The possible involvement of Ca^2+^ binding proteins’ (CaBPs) sequestration of Ca^2+^ could explain this observation and would be an interesting focus for future studies. CaBPs, consisting of chaperones and buffers located throughout the lumen of the ER, help ensure that the [Ca^2+^]_ER_ remains within an appropriate range, which is essential for the maintenance of persistent Ca^2+^ signals and post-translational processing, folding, and export of other proteins (reviewed in Berridge, 2002; Booth and Koch, 1989; Nigam et al., 1994). In the broader setting of virus infection, it is also possible that the E protein could exert an anti-apoptotic activity, as it would post-infection. Upon virus entry, the cellular apoptotic pathway is immediately triggered as a defense mechanism in response to infection. A sudden increase in cytosolic Ca^2+^ could also trigger this pathway, as can the overloading of the mitochondria with Ca^2+^, resulting in a release of cytochrome c and activation of caspase 9, committing the cell to apoptosis (D’Agostino et al., 2005). In this light, it is plausible that the virus hijacks the cell’s Ca^2+^ homeostasis machinery to quickly sequester or export Ca^2+^ from the cytoplasm, in effect, keeping mitochondrial Ca^2+^ levels low to promote cell survival (Pinton et al., 2000; reviewed in Zhou et al., 2009).

Our electrophysiology experiments were designed as an alternative metric for ionotropic E-protein activity. However, unlike the Ca2+ assays, these measurements relied on overexpression of the viral protein on the PM. In this context, electrophysiological currents could not be conclusively distinguished from endogenous or background effects, highlighting persistent ambiguities in E-protein ion-transport function. Earlier reports using truncated SARS-CoV E-protein peptides, reconstituted in artificial lipid bilayers, indicated contrasting and variable permeabilities for Na^+^, K^+^, and Cl^−^ (Regla-Nava et al., 2015; Wilson et al., 2006, 2004). The composition of the bilayer has also been reported to influence E-protein selectivity, with greater cation selectivity in the presence of negatively charged lipid headgroups (Verdiá-Báguena et al., 2012). However, selectivity has often been based on reversal potential measurements confounded by small, variable currents; other electrophysiological characterizations reported poor signal-to-noise profiles with often indistinct gating events (McClenaghan et al., 2020; Wilson et al., 2004, 2006). Considerations such as the regulation of endogenous channels in heterologous expression systems, reproducibility of robust electrical activity, and accurate identification of foreign proteins in reconstituted environments may be particularly important in documenting conduction properties of small viroporins such as the E protein.

Since the [Ca^2+^] gradient across the cytoplasm to the lumen of the ER/Golgi apparatus is one of the highest and most regulated ion gradients observed in cells (reviewed in Berridge et al., 2003; reviewed in Bootman et al., 2001; reviewed in Zhou et al., 2009), one possible explanation for our observations in transfected cells could be explained by the transmembrane voltage that would open the pore and allow Ca^2+^ to permeate through, which is consistent with the 450 mV transmembrane voltage applied in computational electrophysiology simulations by Cao et al (Cao et al., 2020). Consistent with this proposal, our most stable open simulation was produced in the presence of an electric field. Elucidating the effects of transmembrane voltage induced by viroporins would be an interesting line of investigation as these characteristics have been observed for other viruses as well (Clarke et al., 2006; Mehnert et al., 2008; Schubert et al., 1996).

### Limited stability and permeation in reported structures

The quality and usefulness of simulations are predicated on the quality of the initial structure. Our atomistic MD simulations were initiated from the only available E-protein structures, or models based on them, obtained via solution or solid-state NMR. Protein NMR structures can be influenced by modelling assumptions, such as oligomerization state (Cui et al., 2005). Notably, the C-terminal domain of the E protein has been reported as partially disordered (Gadhave et al., 2020), making it challenging to characterize by structural or simulation methods despite its apparent influence on the protein’s cone-like shape and membrane-curvature effects. Given that flexible proteins and ion interactions may be variably described by current protein force fields, we tested a range of simulation conditions and structure variants in pursuit of a stable open model (Table S1). Simulations of the SARS-CoV-2 E-protein structure from solid-state NMR (PDB ID: 7K3G) were incompatible with conduction, and failed to maintain a the 3_10_-helix twist required to satisfy modeling assumptions.

Only one condition, based on solution NMR (PDB ID: 5×29), produced a relatively stable open state in our hands, with a putative hydrophobic gate at the F26-midpoint of the pore. Even this model was stable only on the hundreds-of-nanoseconds time scale, and was not evidently permeable to Ca^2+^. Although several articles and preprints in the past year (Borkotoky & Banerjee, 2020; Cao et al., 2020; Gadhave et al., 2020; Gentile et al., 2020; Gupta et al., 2020; Kuzmin et al., 2021; Yu et al., 2021) have reported simulations of this structure, a review of their results shows similar limitations, with abbreviated timescales (Borkotoky & Banerjee, 2020; Gentile et al., 2020; Gupta et al., 2020), reliance on secondary-structure restraints (Cao et al., 2020), elevated RMSDs (Borkotoky & Banerjee, 2020; Cao et al., 2020), and/or dewetting in the absence of electric fields (Cao et al., 2020). Instability of the open structure may reflect underdetermination of the starting protein or membrane models, unresolved interactions with the C-terminal or other domains, or other factors yet to be identified.

## Conclusion

Taken together, we have shown that recombinantly expressed full-length E protein from SARS-CoV-2 is capable of independent multimerization. We also confirmed that the protein localizes intracellularly, similar to its predecessor from SARS-CoV, with no evidence of ion channel properties at the cell surface. Our coarse-grained simulations further support a role for the E protein in viral budding, as the presence of the protein bends the surrounding membrane. Reduction of intracellular Ca^2+^ in E-protein-transfected cells may further promote viral replication. However, our atomistic simulations and permeation calculations based on previously reported NMR structures resulted in unstable proteins incapable of Ca^2+^ conduction. We emphasize the importance of using high-resolution structural data obtained from a full-length protein to gain detailed molecular insight of the E protein, and enable future drug-screening efforts.

## Data Availability

Input files and representative frames from CG, atomistic MD, and AWH simulations are available on Zenodo (10.5281/zenodo.4818292).

## Acknowledgements

We are grateful for conceptual assistance from Professors Elisa Fadda (Maynooth University) and Ann McDermott (Columbia University), for technical support from Professor Thomas Voets, Marijke Brams and Sara Kerselaers (KU Leuven) and Professor Jan Tytgat (KU Leuven) for earlier experiments on Xenopus oocytes. Financial support for this project was provided by grants from FWO-Vlaanderen (G0C9717N, G0C1319N) and KU Leuven (C3/19/023, C14/17/093), the Knut and Alice Wallenberg foundation (KAW COVID-19 research grant via SciLifeLab), the Swedish Research Council (VR), the Gustafsson Foundation, and the Swedish e-Science Research Center (SeRC). Computational time was provided by the Swedish National Infrastructure for Computing (SNIC), and by Folding@home; we are extremely grateful to all the citizen scientists who contributed their compute power to make this work possible, and to members of the Folding@home community who volunteered to help.

## Author Contributions

Conceptualisation: AM, SPC, YZ, AE, RJH, CU, LD; methodology: AM, SPC, YZ, AE, DP, RJH, CU, LD; software: SPC, YZ, AE, EL, LD; validation: AM, SPC, YZ, AE; formal analysis: AM, SPC, YZ, AE; investigation: AM, SPC, YZ, AE, DP; resources: EL, RJH, CU, LD; data curation: SPC, YZ, AE; original draft: AM, SPC, YZ, AE; review & editing: AM, SPC, YZ, AE, RJH, CU, LD; visualization: AM, SPC, YZ, AE; supervision: RJH, CU, LD; project administration: RJH, CU, LD; funding acquisition: EL, CU, LD.

## Notes

https://zenodo.org/record/4818292#.YLEU8pMzY-Q

### Competing Interest Statement

The authors have declared no competing interest.

