## Supplementary Information for "Probing effects of the SARS-CoV-2 E protein on membrane curvature and intracellular calcium"

### Supplementary Figures

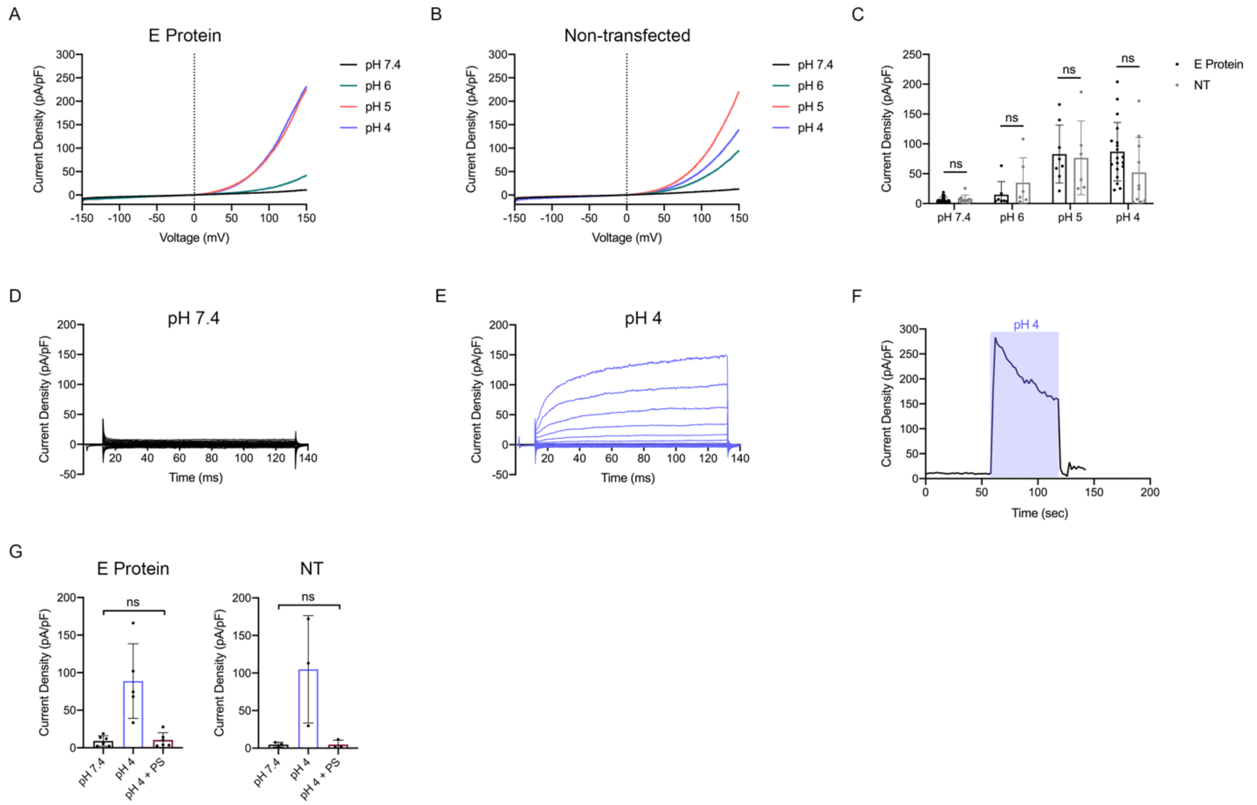


**Figure S1. Evidence of pH-sensitive activity at the plasma membrane in HEK-293 cells.** Cells transiently transfected with the EcGFP construct displayed outward currents in acidic conditions, but could not be significantly distinguished from non-transfected cells. A: Mean traces of EcGFP-transfected cells perfused with standard extracellular solution in decreasing pH, 1 min exposures (pH 7.4 N = 24; pH 6 N = 7; pH 5 N = 8; pH 4 N = 19). B: Mean traces of non-transfected cells under the same conditions as in A (pH 7.4 N = 10; pH 6 N = 6; pH 5 N = 6; pH 4 N = 10). C: Comparison of outward current densities analyzed at +100 mV. Statistical significance was calculated via an unpaired, non-parametric Mann-Whitney test; ns = not significant ($p > 0.05$). D-E: Current traces in response to voltage steps from -160 mV to +40 mV in 20 mV increments from EcGFP-transfected cells. F: Current response in extracellular solution of pH 7.4 vs. F: Time course illustrating on/off kinetics in response to a 1 min exposure of extracellular solution at pH 4 (blue shaded region). G: In a subset of experiments, 50 µM PS was added after switching to an extracellular solution of pH 4, resulting in a nearly complete block of outward current in response to pH. Values in this condition were statistically not significant compared to values at pH 7.4: E protein (N = 6; $p > 0.05$); NT (N = 3; $p > 0.05$); calculated using a paired, non-parametric Wilcoxon test.

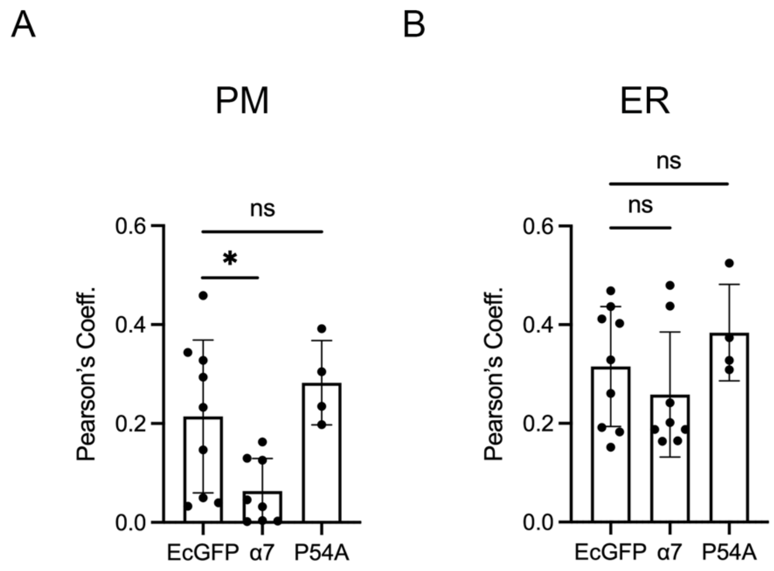


**Figure S2. Modification of signal and retention motifs do not increase expression to plasma membrane.**

Pearson's coefficients were calculated using replicate confocal images to determine coexpression of the WT E protein (EcGFP), the E protein with N-terminal α7 export signal, or the E protein with a point mutation at P54 with signals corresponding to the PM (A) or ER (B). A: Pearson's coefficient ($r$) for EcGFP signal with PM signal = 0.22 ± 0.21; $r_{\alpha 7:PM}$ = 0.063 ± 0.066; $r_{P54A:PM}$ = -0.29 ± 0.085. B: $r_{EcGFP:ER}$ 0.32 ± 0.13; $r_{\alpha 7:ER}$ = 0.26 ± 0.13; $r_{P54A:ER}$ = 0.38 ± 0.98. Statistical significance was calculated via a paired two-tailed $t$-test between WT and modified signal (* $p < 0.05$; ns = not significant).

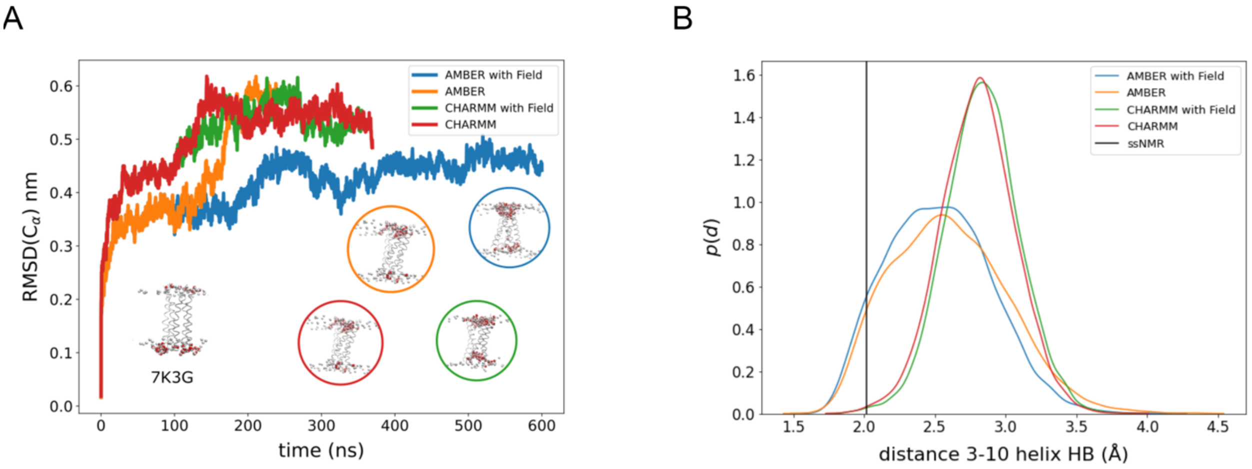


**Figure S3. Limited stability of the closed solid-state NMR structure (PDB ID 7K3G)**
A: Root mean-squared deviations (RMSDs) of Cα atoms as a function of time during simulations of the pentameric solid-state NMR structure (PDB ID: 7K3G). Models were simulated in the CHARMM36m (red, green) or AMBER99SB-ILDN + SLIPIDS (blue, orange) force fields in the absence (red, orange) or presence (green, blue) of an electric field. Inset snapshot at lower left shows the starting model, viewed from membrane plane; additional insets show snapshots from the end of the simulations. B: Probability distribution of the backbone hydrogen-bond distance (O of F20 to H of F23) that defines the $3_{10}$-helix transition, in the four simulations colored as in *A*. The vertical black line indicates the hydrogen-bond distance in the solid-state NMR structure.

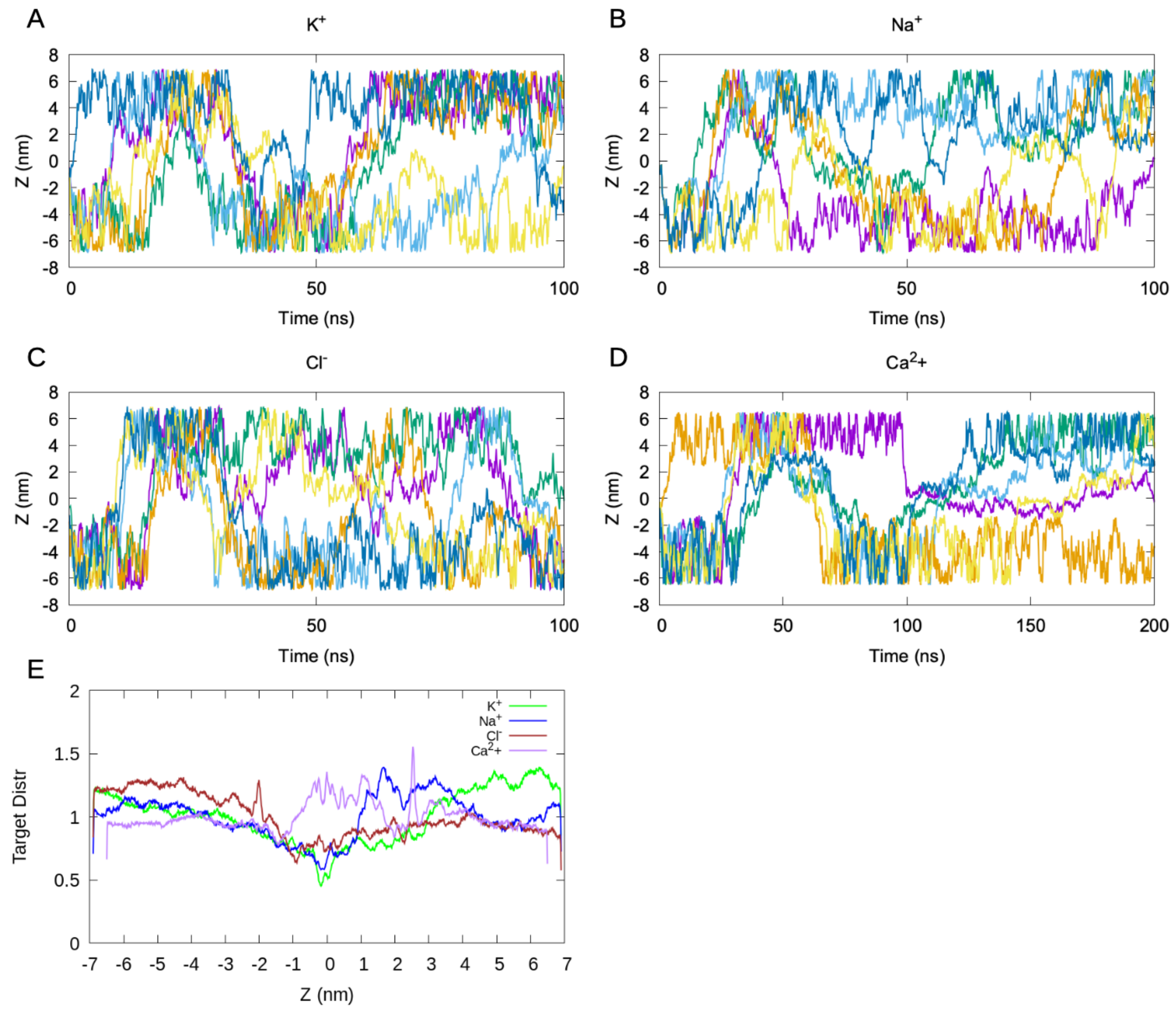


**Figure S4: Convergence of ion-permeation simulations in the most stable E-protein simulation model.**

Ion movement along the Z-axis of the pore as a function of time in accelerated-weight histogram simulations of permeation through the most stable E-protein simulation model by A: K+, B: $Na^+$, C: $Cl^-$, or D: $Ca^{2+}$. Different walkers are shown in different colors. E: Target distribution as a function of the pore Z-axis for $K^+$ (green), $Na^+$ (blue), $Cl^-$ (red) and $Ca^{2+}$ (purple) ion-permeation simulations.

**Table S1: Atomistic MD simulations.** Conditions are identified by starting structures (PDB ID and ensemble-model number; pentamerized monomers, 'x 5'; homology model based on SARS-CoV variant, 'mod.'), post-translational modifications (PTM; palmitoylated, 'palm.'; glycosylated, 'glyc.'), equilibration protocols (CHARMM-GUI default protocol, 'GUI'; with an electric field, '+ Field'; with pore restraints based on Ref. 38, '+ Pore restraints'), and production conditions (CHARMM36m force field, 'CHARMM'; AMBER99SB-ILDN + SLipids force field, 'AMBER'). Most simulations produced asymmetrized (asymm.) or collapsed endpoints, with only simulation 8 producing a possibly conductive state over hundreds of nanoseconds.

| ID | Structure | PTM | Equilibration | Production | Result |
|---|---|---|---|---|---|
| 1 | 2MM4.1 x 5, mod. | None | GUI | CHARMM | Asymm., collapsed |
| 2 | 2MM4.1 x 5, mod. | None | + Field | CHARMM + Field | Asymm., collapsed |
| 3 | 2MM4.1 x 5, mod. | Palm. | + Field | CHARMM | Asymm., collapsed |
| 4 | 2MM4.1 x 5, mod. | Palm. | + Field | CHARMM + Field | Asymm., collapsed |
| 5 | 5X29.1, mod. | None | GUI | CHARMM | Asymm., collapsed |
| 6 | 5X29.1, mod. | None | GUI | CHARMM | Mostly symmetric, collapsed |
| 7 | 5X29.1, mod. | None | + Pore restraints | CHARMM | Asymm., collapsed |
| 8 | 5X29.1, mod. | None | + Pore restraints | CHARMM + Field | Mostly symmetric, possibly open |
| 9 | 5X29.1, mod. | None | GUI | CHARMM + Field | Mostly symmetric, collapsed |
| 10 | 5X29.1, mod. | Palm. | GUI | CHARMM | Asymm., collapsed |
| 11 | 5X29.1, mod. | Palm., glyc. | GUI | CHARMM | Asymm., collapsed |
| 12 | 5X29.1, mod. | Glyc. | GUI | CHARMM | Asymm., collapsed |
| 13 | 5X29.6, mod. | None | + Pore restraints | CHARMM + Field | Asymm., collapsed |
| 14 | 5X29.7, mod. | None | + Pore restraints | CHARMM + Field | Asymm., collapsed |
| 15 | 7K3G.1 | None | + Pore restraints | AMBER + Field | Asymm., collapsed |
| 16 | 7K3G.1 | None | + Pore restraints | AMBER | Mostly symmetric, collapsed |
| 17 | 7K3G.1 | None | + Pore restraints | CHARMM + Field | Mostly symmetric, Collapsed |
| 18 | 7K3G.1 | None | + Pore restraints | CHARMM | Mostly symmetric, collapsed |
